# Clustering Strategies Improve Structure-Preserving Visualization of Single-Cell RNA-seq Data with CBMAP

**DOI:** 10.64898/2026.04.30.721861

**Authors:** Meysa Alchaar, Berat Doğan

## Abstract

Dimensionality reduction for visualization is a fundamental step in single-cell RNA sequencing (scRNA-seq) analysis due to the extremely high dimensionality of gene expression profiles. However, widely used nonlinear embedding techniques such as UMAP and t-SNE can introduce substantial distortions when projecting data into two-dimensional space, potentially altering global organization, local neighborhoods, and distance relationships in ways that may mislead downstream biological interpretation. In this study, we investigate the applicability of Clustering-Based Manifold Approximation and Projection (CBMAP) for the visualization of scRNA-seq data and systematically examine how clustering strategies influence the quality of the resulting embeddings. CBMAP was integrated with several clustering algorithms commonly used in single-cell analysis, including k-means, Leiden, HDBSCAN, Secuer, HGC, and FlowSOM. The resulting embeddings were evaluated using quantitative metrics that measure global, local, and distance-level structure preservation and were compared with widely used dimensionality reduction methods such as UMAP, t-SNE, and PaCMAP across multiple benchmark datasets. Our results demonstrate that the clustering stage plays a critical role in determining the structural fidelity of CBMAP embeddings. Clustering algorithms specifically designed for single-cell transcriptomic data, particularly Secuer, produced more consistent preservation of global relationships between cell populations. Across multiple datasets, CBMAP more faithfully preserved global structural organization and inter-population distance relationships than the compared methods, although local neighborhood preservation was generally weaker than in techniques optimized for local structure. Importantly, CBMAP embeddings retained biologically meaningful relationships in trajectory benchmark datasets. When combined with RNA velocity analysis, CBMAP successfully preserved cyclic progenitor states and branching differentiation trajectories, demonstrating compatibility with trajectory-aware visualization. These findings indicate that CBMAP provides a structure-faithful visualization framework for scRNA-seq data and that clustering selection plays a central role in determining embedding quality.

## 1. Introduction

Single-cell RNA sequencing (scRNA-seq) technologies have become fundamental tools in biological research and precision medicine. Their importance stems from their ability to dissect cellular heterogeneity within tissues and to characterize the diverse functional states of individual cells. By profiling gene expression at single-cell resolution, scRNA-seq enables the investigation of cellular functions, developmental trajectories, and dynamic responses to internal and external influences. Projects such as the Single Cell Atlas (SCA) [1] and the Human Cell Landscape (HCL) [2] have produced large-scale single-cell gene expression datasets from different organs. Despite their scientific value, these datasets are extremely high-dimensional, heterogeneous, and often sparse, introducing numerous computational challenges for downstream analysis [3]. The early stages of scRNA-seq analysis, such as quality control and gene expression quantification, depend on bioinformatics methods that can substantially influence biological interpretations [4]. The resulting gene expression matrices are characterized by sparse count data with a high proportion of zeros, non-Gaussian distributions, and substantial technical noise, often accompanied by irregular clusters [4-5].

One of the most critical stages in the scRNA-seq analysis pipeline is reducing the complexity caused by the extremely large number of measured genes. Principal Component Analysis (PCA) [6] is a widely used linear decomposition method that transforms the original gene expression matrix into a set of orthogonal components. These components capture the directions of greatest variance in the data, allowing the dataset to be summarized into a smaller number of informative features while preserving the most biologically meaningful structure. However, when dimensionality reduction is applied for visualization purposes, the task becomes significantly more challenging. In scRNA-seq analysis, it is common to use UMAP [7] and t-SNE [8], which are nonlinear dimensionality reduction methods designed to represent relationships between cells in low-dimensional space. Nevertheless, numerous studies and benchmarking analyses [9-12] have shown that UMAP, t-SNE, and other nonlinear visualization techniques can introduce substantial distortions when projecting data into two or three dimensions. These distortions may alter global organization, local neighborhood structures, and distance relationships, potentially affecting downstream biological interpretations, as clearly demonstrated in [11].

In this study, we examine a recently published dimensionality reduction method, CBMAP (Clustering-Based Manifold Approximation and Projection) [13], and investigate its applicability to scRNA-seq data. In its original formulation, CBMAP employs k-means clustering to identify representative structures that guide the manifold approximation process.

However, the heterogeneous, sparse, and noise-prone characteristics of scRNA-seq datasets often violate the assumptions underlying k-means clustering and may limit the robustness of the resulting embeddings. To address this limitation, we explore the integration of CBMAP with several clustering algorithms commonly used in single-cell analysis and better suited for capturing complex cellular population structures. Specifically, CBMAP was combined with Leiden, HDBSCAN, Secuer, HGC, and FlowSOM clustering approaches in order to evaluate how clustering behavior influences the structural fidelity of the resulting low-dimensional embeddings. A comprehensive set of supervised and unsupervised evaluation metrics was employed to identify the clustering strategies most appropriate for scRNA-seq datasets. Furthermore, CBMAP embeddings generated with different clustering strategies were systematically compared with widely used nonlinear dimensionality reduction methods to assess their ability to preserve global and local structural relationships in single-cell data. In addition, trajectory benchmark datasets were analyzed using RNA velocity to evaluate whether CBMAP embeddings preserve biologically meaningful developmental dynamics. To our knowledge, this study represents the first systematic investigation of clustering-driven CBMAP embeddings for scRNA-seq visualization.

## 2. Materials and Methods

### 2.1. Datasets

The performance of CBMAP was assessed using multiple publicly available scRNA-seq datasets, summarized in Table 1. These datasets represent diverse biological systems and varying levels of cellular heterogeneity, allowing the evaluation of CBMAP under different data complexity scenarios.

**Table 1.** A description of scRNA-seq datasets used in the experiments.

| **Dataset** | **Tissue** | **Cells** | **Genes** | **Number of Cell Type** |
| --- | --- | --- | --- | --- |
| Duo [14] | PBMCs | 3994 | 15715 | 4 or 8 |
| Muraro [15] | Pancreas | 2282 | 18962 | 9 |
| Kang [16] | PBMCs | 13999 | 14053 | 13 |
| Smart-seq [17] | Mouse ventromedial hypothalamus (VMH) neurons | 3850 | 1999 | 28 |
| Utero E8.5 [18] | Ex and In utero mouse embryo E8.5 | 10290 | 19588 | 19 |

### 2.2. Clustering algorithms

CBMAP is a dimensionality reduction method designed to preserve the structural properties of continuous data during the embedding process. The method first clusters data points in the high-dimensional space and computes the membership of each point with respect to the cluster centers. The cluster centers are then projected into a low-dimensional space using PCA, and each data point is initialized near its corresponding cluster center. The final low-dimensional embedding is obtained by minimizing the discrepancy between high- and low-dimensional memberships using a membership-based objective function defined by the Frobenius norm.

Previous studies have shown that CBMAP effectively preserves both global and local structural relationships in benchmark datasets [13]. In particular, the intrinsic geometry of continuous manifolds can be maintained when embedding data into two-dimensional space, as illustrated in classical examples such as the Swiss-roll and Mammoth datasets, where the underlying shapes and cluster structures remain visually distinguishable. This property is often degraded in other nonlinear dimensionality reduction techniques [10-11].

In its original formulation, CBMAP relies on the k-means algorithm as the clustering step guiding the manifold approximation process. However, the heterogeneous and noise-prone nature of scRNA-seq datasets may limit the effectiveness of k-means clustering. Therefore, in this study we investigate the integration of CBMAP with clustering algorithms commonly used in single-cell analysis in order to evaluate whether more suitable clustering strategies can improve the structural fidelity of CBMAP embeddings for single-cell transcriptomic data.

Clustering is a key step in scRNA-seq analysis, as it separates cells into biologically meaningful subpopulations corresponding to distinct cell types or cellular states. Various approaches have been proposed for clustering single-cell transcriptomic data, including methods derived from k-means, hierarchical, graph-based, and density-based clustering [19], as well as more recent deep learning–based strategies [20]. Comparative studies indicate that the optimal clustering method depends on both the research objective and the characteristics of the dataset [21-23]. In this study, clustering algorithms were selected based on practical considerations relevant to their integration with the CBMAP framework. Specifically, the selected methods were required to accept preprocessed data as input rather than performing internal preprocessing steps, unlike approaches such as SC3 [24], while also providing straightforward implementation, computational efficiency, and low sensitivity to user-defined parameters. Five clustering algorithms satisfying these criteria and demonstrating favorable performance in preliminary CBMAP experiments were selected for further analysis (Table 2).

**Table 2.** A description of clustering algorithms used in CBMAP for scRNA-seq analysis.

| Method | Type | Framework | Parameters |
| --- | --- | --- | --- |
| Leiden [25] | Graph-based clustering | Uses a graph-based community detection framework that optimizes modularity on a k-nearest neighbor graph | Resolution parameter |
| HDBSCAN [26] | Density-based clustering | Hierarchical density-based spatial clustering of applications with noise that find clusters of varying densities | Minimum cluster size (20 for all experiments) |
| Secuer [27] | Spectral clustering | Uses an anchor-based spectral clustering that represents cells via a reduced cell-anchor graph | No parameter required |
| HGC [28] | Graph-based hierarchical clustering | Hierarchical Graph-based Clustering that constructs a dendrogram of cells on their shared nearest-neighbor (SNN) graph. | Number of clusters and neighbors |
| FlowSOM [29] | Model-based | Uses a self-organizing map (SOM) to project high-dimensional cytometry data onto a low-dimensional grid, followed by meta-clustering of SOM nodes | Number of clusters and grid size (7 for all experiments) |

During preliminary experiments on scRNA-seq datasets, we observed that CBMAP is sensitive to outliers, even when these correspond to biologically meaningful rare cell populations. To improve robustness, cluster centers in the CBMAP framework were computed using the median of cluster members instead of the mean used in the original algorithm, allowing the centers to better represent the underlying data distribution in the presence of extreme values.

In addition, a distance-based outlier detection strategy based on the three-sigma (3σ) rule was applied in the PCA-transformed space of the data. Cells whose distances exceeded the threshold exceeding *μ*_*d*_ *+ kσ*_*d*_ were identified as potential outliers, where *μ*_*d*_ denotes the mean distance, *σ*_*d*_ represents the standard deviation of the distances, and *k* ∈ {2,3}. This procedure was applied to all clustering algorithms evaluated in this study except HDBSCAN, which internally identifies outliers and labels them as −1.

Detected outlier cells were subsequently assigned to clusters using one of two strategies. In the first approach, k-means clustering was applied to the outlier set with the number of clusters selected from the range 2-5 based on silhouette scores. In the second approach, a k-nearest neighbors (kNN) classifier trained on the inlier cells was used to assign each outlier cell to the cluster of its nearest neighbors.

### 2.3. Structural preservation metrics for comparing clustering strategies in CBMAP

To evaluate the influence of different clustering algorithms on CBMAP embeddings, we quantified the ability of the resulting low-dimensional representations to preserve both global and local structural properties of the original high-dimensional data. Two complementary metrics were used for this purpose.

Global structure preservation was assessed using the Random Triplet Accuracy (RTA) metric. RTA evaluates whether relative distance relationships among randomly selected triplets of data points are preserved after dimensionality reduction. For each triplet (i, j, k), the metric verifies whether the ordering of pairwise distances in the original space is maintained in the embedded space. Higher RTA values indicate better preservation of global geometric relationships.

Local structure preservation was measured using the Nearest Neighbor Kept (NNKept) metric, which quantifies the proportion of nearest-neighbor relationships retained after dimensionality reduction. For each cell, the nearest neighbors identified in the high-dimensional space are compared with those obtained in the low-dimensional embedding, and the fraction of preserved neighbors is reported as the NNKept score.

To facilitate objective comparison of clustering strategies, CBMAP provides the try_clustering function, which evaluates embedding quality under different scoring modes. Preservation can be assessed locally using NNKept, globally using RTA, or through a combined score defined as 0.5 RTA+0.5 (10×NNKept), Since NNKept values typically fall within the range 0–0.2 while RTA ranges between 0–1, the NNKept component is scaled by a factor of 10 to ensure balanced contribution of both metrics in the combined score.

### 2.4. Evaluation metrics for dimensionality reduction methods

To provide a comparative perspective, CBMAP embeddings were compared with those generated by widely used nonlinear dimensionality reduction methods, including UMAP, t-SNE, and PaCMAP. PaCMAP is a nonlinear manifold learning method designed to better preserve global data structure while maintaining local neighborhood relationships [30]. This comparison allows the systematic evaluation of CBMAP’s ability to preserve both global organization and local neighborhood structures in single-cell transcriptomic data.

Dimensionality reduction methods are frequently used without systematic evaluation of the reliability of their embeddings, which may lead to distorted or misleading representations of high-dimensional biological data. In addition, many dimensionality reduction techniques are sensitive to user-defined parameters, potentially introducing biased biological interpretations [11-12]. To address these concerns, a set of evaluation metrics was employed to assess the structural properties of the generated embeddings.

An evaluation framework for dimensionality reduction methods in biological applications was introduced in [10], which proposed quantitative measures for evaluating global, local, and distance preservation properties of embeddings. In addition, we adopted selected metrics from [11], which analyzed distortions introduced by commonly used visualization techniques such as UMAP and t-SNE in scRNA-seq datasets. Based on these studies, embedding quality was evaluated at three complementary levels: global structure preservation, local neighborhood preservation, and distance relationship preservation.

#### 2.4.1. Global structure preservation

Global structure preservation reflects the ability of an embedding to maintain large-scale relationships between cellular populations. To quantify global preservation, Random Triplet Accuracy (RTA) was used, which measures the proportion of randomly sampled point triplets for which the relative ordering of pairwise distances is preserved between the high-dimensional and low-dimensional spaces.

As an additional unsupervised metric, distance Spearman correlation was employed, measuring the Spearman rank-order correlation between pairwise distances in the high-dimensional and low-dimensional spaces. Because computing all pairwise distances can be computationally expensive for large scRNA-seq datasets, these metrics were calculated on randomly sampled subsets of the data.

Global relationships were also evaluated at the cluster level by examining the preservation of k-nearest neighbor relationships among cluster centroids. In addition, centroid distance correlation was computed as the Spearman correlation between centroid distance matrices in the high- and low-dimensional spaces. Another global-level evaluation approach assesses preservation at the cell-type level using Kendall’s Tau correlation, which compares the ranking of neighboring cell types in the high-dimensional space with their ranking in the embedding.

#### 2.4.2. Local structure preservation

Local structure preservation evaluates whether neighborhood relationships among cells are maintained after dimensionality reduction. In this study, an unsupervised neighbor preservation metric was employed. For each point i, the set of k=5 nearest neighbors in the high-dimensional space N(i) was compared with the corresponding neighbor set N′(i) in the low-dimensional embedding. The average proportion of preserved neighbors across all points was then calculated.

In addition, the Jaccard distance metric was also applied. In this approach, the 30 nearest neighbors of each cell are identified in both the high- and low-dimensional spaces, and the dissimilarity between the two neighbor sets is measured using the Jaccard distance.

#### 2.4.3. Distance relationship preservation

Distance preservation was evaluated using the strategy proposed in [11], which analyzes groups of cells with well-defined distance relationships in the high-dimensional space, including sets of approximately equidistant cells. By examining how these relationships are represented in the embedding, it is possible to assess the extent to which relative distance patterns are maintained after dimensionality reduction.

Additionally, the degree of separation between cell types was evaluated by comparing the distributions of intra-type and inter-type pairwise distances. The difference between these distributions was quantified using the two-sample Kolmogorov–Smirnov (K–S) statistic.

## 3. Results and Discussion

### 3.1. Effect of clustering strategy on CBMAP embedding quality

To investigate how clustering strategies influence CBMAP embeddings, we evaluated the selected clustering algorithms using the structural preservation metrics described in Section 2.3. The results for the Muraro dataset are summarized in Fig. 1, where global structure preservation is assessed using Random Triplet Accuracy (RTA) and local structure preservation using the Nearest Neighbor Kept (NNKept) metric. Additional results for the Duo 4Eq, Kang, SMART-seq Mouse, and Utero E8.5 Embryo datasets are provided in Supplementary Fig. S1.

**Figure 1.**
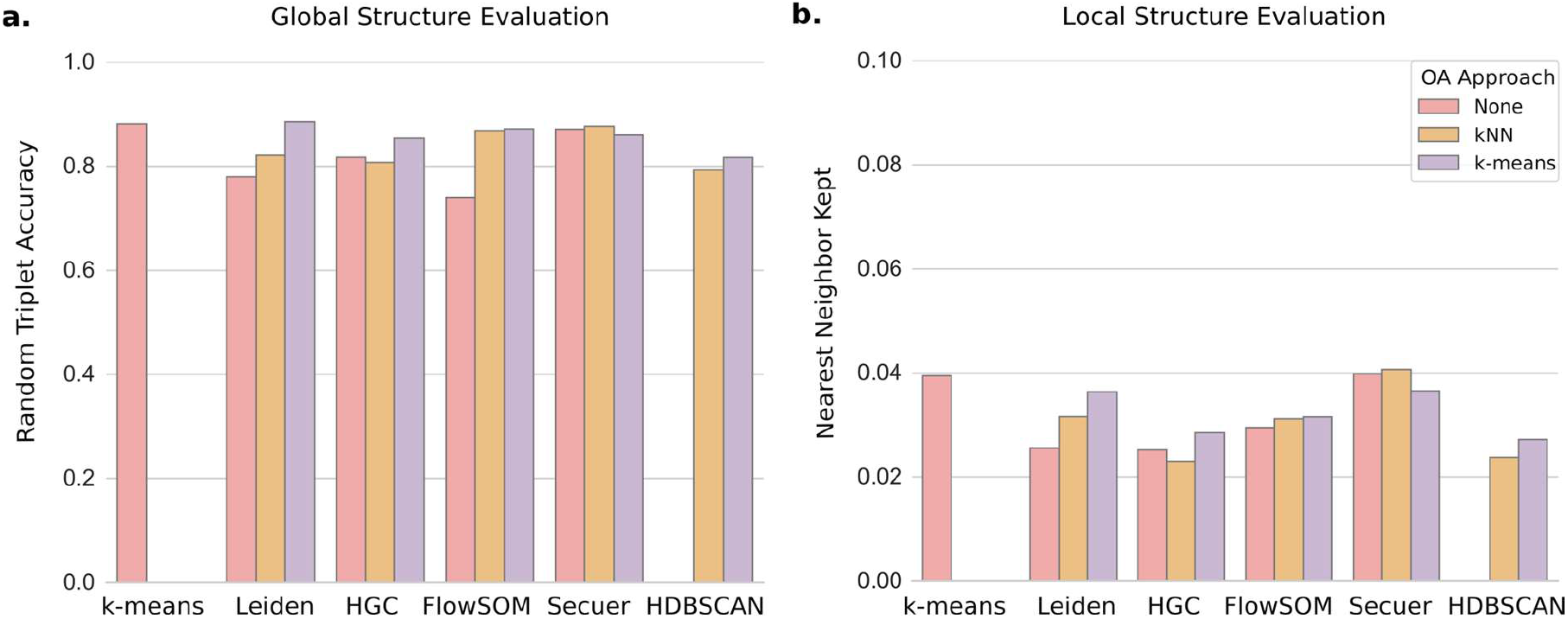
CBMAP embeddings generated using different clustering algorithms on the Muraro dataset. Bar plots show the average evaluation scores across five runs for each clustering method. Unless otherwise specified, default parameters were used (number of clusters = 27, corresponding to three times the number of annotated cell types, for k-means, HGC, and FlowSOM; resolution = 1.7 for Leiden; number of neighbors = 20 for HGC). Three outlier-handling strategies were evaluated: no outlier handling, sigma-rule–based outlier detection followed by assignment to the nearest inlier cluster using k-nearest neighbors (kNN), and clustering of detected outliers using k-means. **(a)** Global structure preservation evaluated using the Random Triplet Accuracy (RTA) metric. **(b)** Local structure preservation evaluated using the Nearest Neighbor Kept (NNKept) metric.

The results show that the quality of CBMAP embeddings is strongly dependent on the clustering strategy used in the initial stage of the algorithm. Although the original CBMAP implementation relies on k-means clustering, several alternative clustering methods produced embeddings with improved structural preservation. On the Muraro dataset, k-means yielded strong global preservation but was not consistently optimal once alternative clustering strategies and outlier-aware approaches were considered. In particular, Leiden, FlowSOM, and Secuer frequently achieved equal or higher RTA values, indicating improved preservation of global relationships between cellular populations. A similar dependence was observed for local structure preservation. As shown in Fig. 1b, the clustering strategy substantially affected NNKept scores, with Secuer providing the strongest overall local preservation on the Muraro dataset and Leiden also benefiting noticeably from outlier-aware processing. These findings suggest that clustering methods that better capture the intrinsic organization of scRNA-seq data tend to provide a more suitable initialization for CBMAP. At the same time, the effect of outlier handling was not uniform across all methods. For some clustering algorithms, especially Leiden, FlowSOM, and HDBSCAN, outlier-aware strategies improved both global and local preservation, whereas for others the gains were smaller or more selective. This pattern indicates that robustness to rare or noisy cell populations is an important but method-dependent factor in CBMAP performance.

Among the evaluated methods, Secuer emerged as the most balanced clustering strategy for CBMAP. It consistently produced competitive or superior results across both global and local preservation metrics, while requiring relatively limited parameter tuning. This behavior is also evident in Fig. 2a, where the Kang dataset embedding generated with Secuer shows clearer recovery of specific populations, particularly the Mk cells, than the corresponding k-means-based embedding. On the Muraro dataset, both HGC and Leiden produced interpretable embeddings (Fig. 2b), but with a different trade-off: HGC showed slightly stronger global preservation, whereas Leiden achieved slightly better local preservation. This again highlights that the clustering stage influences not only overall embedding quality but also the balance between local and global structure retention.

**Figure 2.**
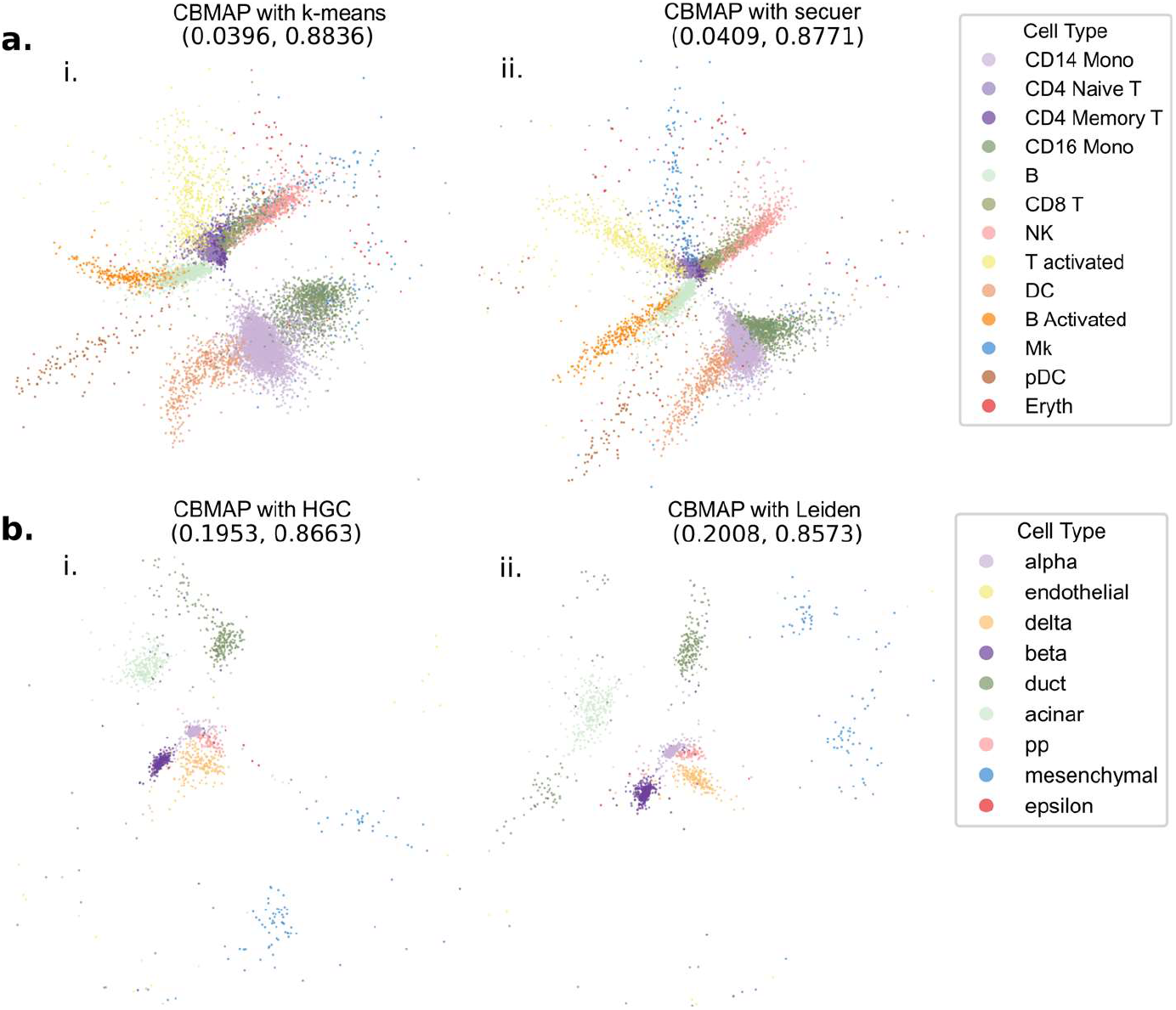
CBMAP visualizations generated using different clustering strategies. **(a)** CBMAP embeddings of the Kang dataset obtained using (i) k-means clustering with the number of clusters set to 39 (three times the number of annotated cell types) and (ii) the Secuer clustering algorithm with default parameters. **(b)** CBMAP embeddings of the Muraro dataset obtained using (i) HGC clustering with 39 clusters and 20 nearest neighbors and (ii) Leiden clustering with a resolution parameter of 1.7. The NNKept and RTA values, representing local and global structure preservation, are reported in parentheses.

Importantly, the trends observed in the Muraro dataset were not dataset-specific. Supplementary Fig. S1 shows that clustering sensitivity is reproducible across additional scRNA-seq datasets, although the best-performing method varies somewhat with dataset structure. This result is consistent with the design of CBMAP, which uses cluster structure to guide the manifold approximation process. Consequently, inaccuracies or mismatches introduced at the clustering stage can propagate directly into the final low-dimensional representation.

Overall, these findings demonstrate that clustering is not merely a preprocessing step for CBMAP, but a central determinant of embedding quality. Careful selection of clustering strategies, particularly those developed for single-cell transcriptomic data, can substantially improve the structural fidelity and interpretability of CBMAP visualizations.

### 3.2. Comparison of CBMAP with existing dimensionality reduction methods

To further assess the behavior of CBMAP, we compared its embeddings with those produced by widely used nonlinear dimensionality reduction methods, including UMAP, t-SNE, and PaCMAP. Representative visual comparisons for the Muraro dataset are shown in Fig. 3, while quantitative evaluations based on global, local, and distance-related structure preservation are presented in Figs. 4–7 and Supplementary Figs. S2–S7.

**Figure 3.**
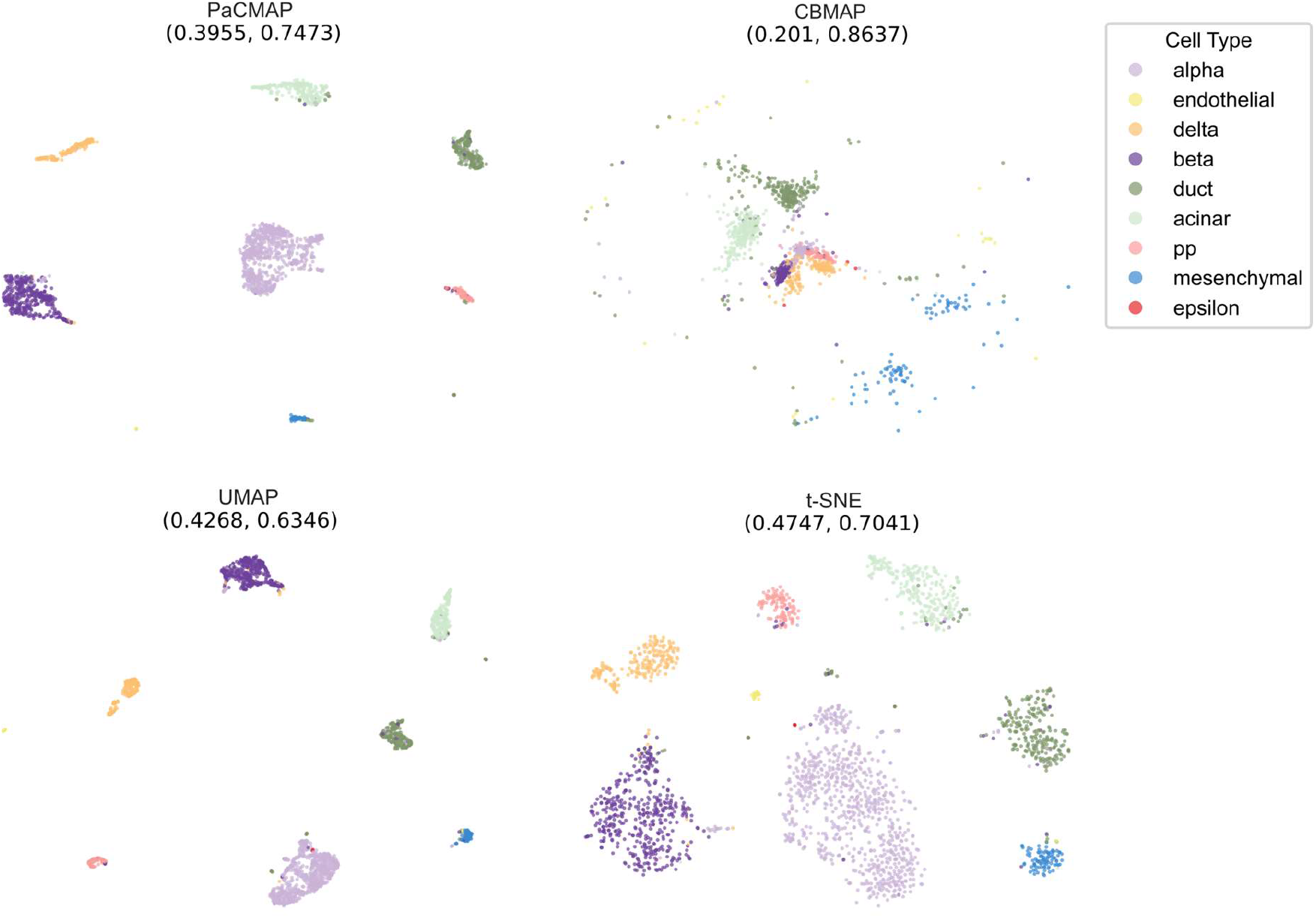
Comparison of dimensionality reduction methods on the Muraro dataset. Embeddings generated using CBMAP, UMAP, t-SNE, and PaCMAP are shown. All methods were implemented using their default parameter settings. For CBMAP, the HGC clustering algorithm was used to generate the embeddings. The values reported in parentheses correspond to the NNKept and RTA scores, representing local and global structure preservation, respectively.

**Figure 4.**
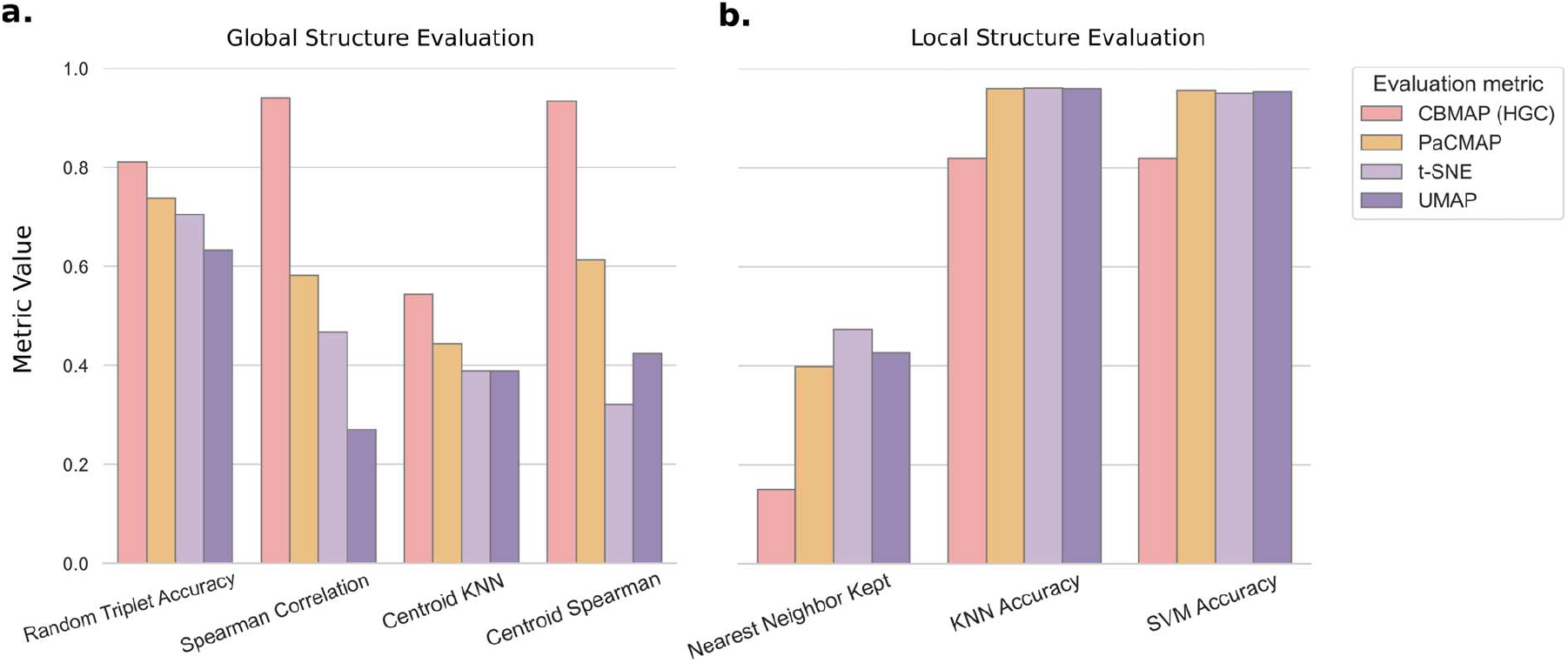
Evaluation of structure preservation for different dimensionality reduction methods on the Muraro dataset. Bars represent the mean scores across five runs. **(a)** Global structure preservation metrics. **(b)** Local structure preservation metrics.

Visual inspection already reveals a clear difference in embedding behavior. UMAP, t-SNE, and PaCMAP tend to generate compact, strongly separated island-like clusters. In contrast, CBMAP produces a more continuous layout in which multiple populations remain positioned relative to one another. CBMAP retains a relational organization that more closely reflects continuity and between-population positioning. This visual pattern is consistent with the design objective of CBMAP, which prioritizes structural preservation over maximal visual separation.

#### 3.2.1. Global structure preservation

Quantitative evaluation strongly favors CBMAP for global structure preservation. As shown in Fig. 4a, CBMAP achieved the highest scores on all evaluated global metrics in the Muraro dataset, including Random Triplet Accuracy, distance Spearman correlation, centroid kNN preservation, and centroid Spearman correlation. These results indicate that CBMAP preserves large-scale geometric relationships among cells more faithfully than UMAP, t-SNE, and PaCMAP. This advantage is not limited to the Muraro dataset. Supplementary Fig. S4 shows the same overall tendency across Duo 4Eq, Kang, SMART-seq Mouse, and Utero E8.5 Embryo, demonstrating that strong global preservation is a recurrent property of CBMAP rather than an isolated result. The same conclusion is supported by the cell-type-level analysis in Fig. 5a and Supplementary Fig. S5. Using Kendall’s Tau correlation of cell-type neighbor rankings relative to the PCA reference space, CBMAP consistently produced higher correlations than the competing methods. Thus, CBMAP better preserves the relative proximity structure between cell populations, not only at the level of individual cells but also at the level of biologically meaningful cell types. From a biological perspective, this is a major advantage. In scRNA-seq data, the arrangement of populations relative to one another often carries important information about lineage relationships, intermediate states, or partial continuity between cell identities. By preserving this organization more faithfully, CBMAP reduces the risk of interpreting visualization artifacts as biological discreteness.

**Figure 5.**
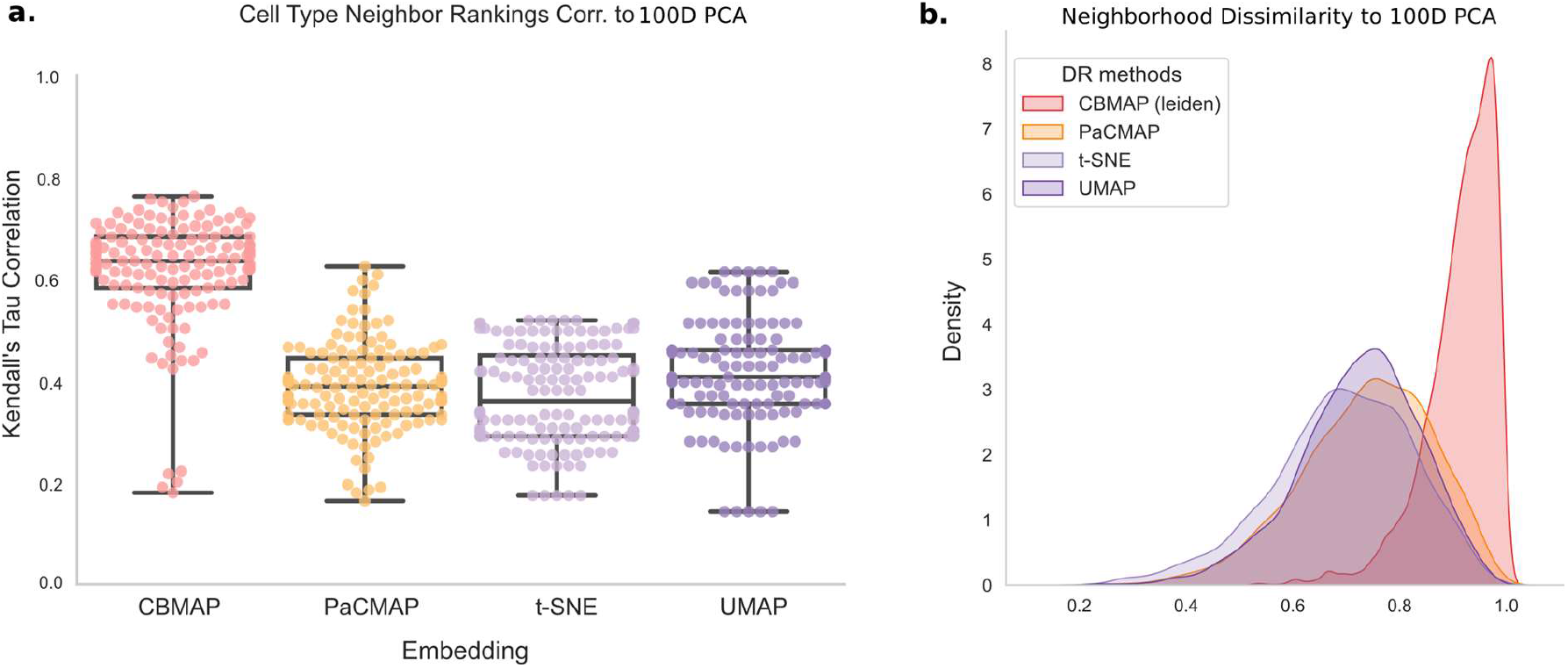
Evaluation of structure preservation for different dimensionality reduction methods on the SMART-seq Mouse dataset. **(a)** Box plots showing the correlations between cell-type neighbor rankings in the two-dimensional embeddings and those in the higher-dimensional PCA space. Each embedding method was repeated five times; higher correlations indicate better preservation of global cell-type relationships. **(b)** Distribution of Jaccard distances between cell neighborhoods in the two-dimensional embeddings and the higher-dimensional PCA space. Lower Jaccard distances indicate better agreement between neighborhood structures.

#### 3.2.2. Local structure preservation

Local structure preservation showed a different pattern. As illustrated in Fig. 4b, CBMAP generally achieved lower local preservation scores than UMAP, t-SNE, and PaCMAP, particularly for NNKept. This indicates that the competing methods are more effective at preserving exact nearest-neighbor relationships in two-dimensional space. The same trend is reinforced by the Jaccard-distance analysis in Fig. 5b and Supplementary Fig. S5, where CBMAP produced higher neighborhood dissimilarity values relative to the PCA reference space. These results confirm that CBMAP does not optimize local neighborhood reconstruction as aggressively as methods such as UMAP and t-SNE. However, the local preservation results should be interpreted in context. First, CBMAP still retained meaningful local structure, as reflected by non-trivial NNKept, kNN, and SVM scores across datasets. Second, the weaker local preservation appears to be a consequence of the same design choice that enables stronger global preservation. Methods such as UMAP and t-SNE explicitly emphasize local neighborhood reconstruction, often at the expense of global geometry. CBMAP instead preserves broader structural organization while maintaining a reasonable, though less optimized, degree of local continuity. This trade-off may be desirable in settings where the primary goal is not to produce the most compact local clusters, but to preserve the relative spatial logic of the original data. In such cases, a modest reduction in nearest-neighbor fidelity may be acceptable if it avoids stronger distortions of global organization.

#### 3.2.3. Distance relationship preservation

Distance-based analyses further clarify the distinct behavior of CBMAP. Using the evaluation framework proposed in [11], we examined equidistant cell groups in the Kang dataset (Fig. 6). In the UMAP embedding, near and far equidistant groups appear visually similar, indicating substantial compression of quantitative distance relationships. In contrast, CBMAP preserves a clearer distinction: near equidistant cells remain relatively compact, whereas far equidistant groups are more dispersed. Although this does not represent perfect metric preservation in two dimensions, it indicates that CBMAP retains relative distance information more effectively than UMAP in this setting.

**Figure 6.**
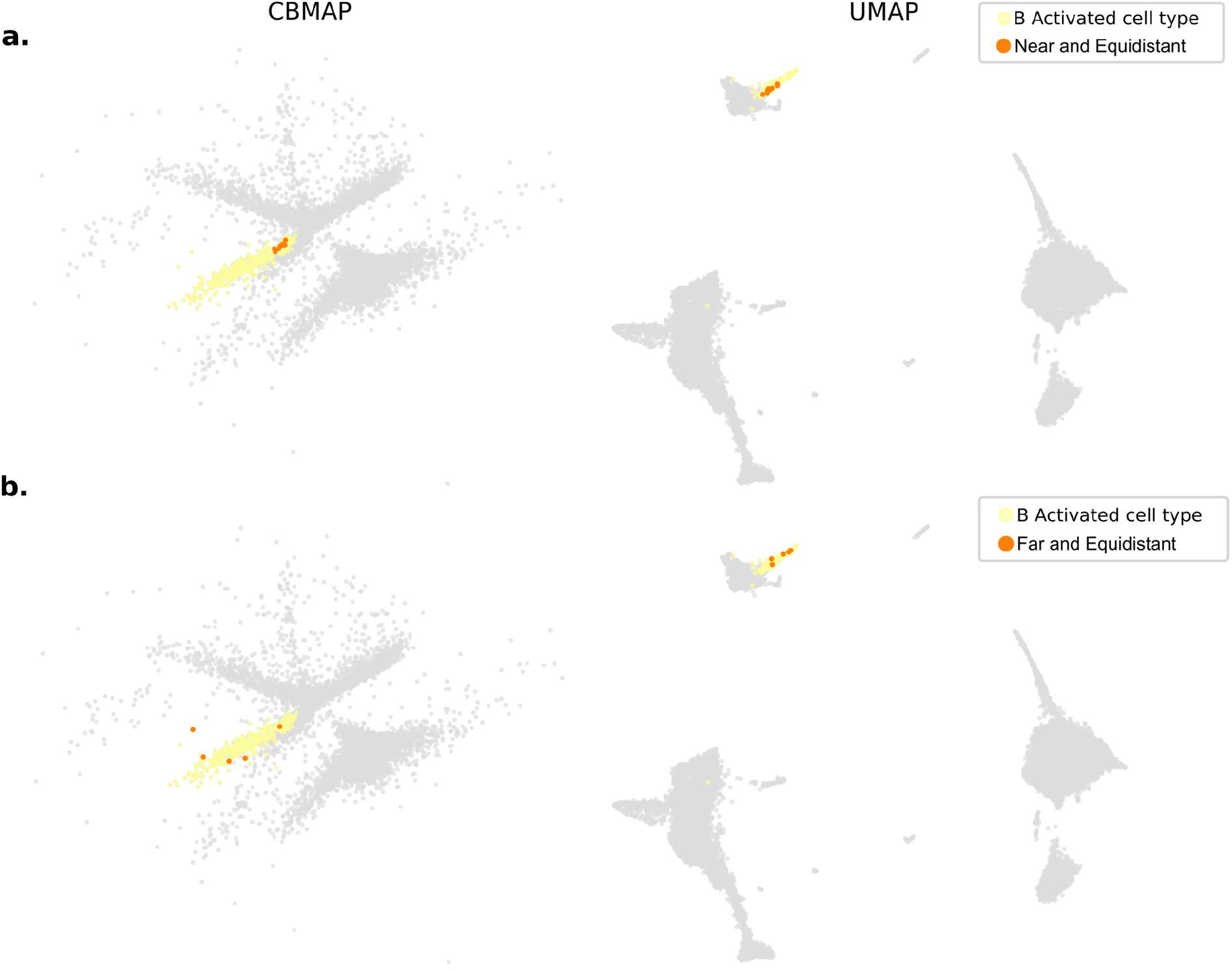
Distance preservation analysis using equidistant cell groups in CBMAP and UMAP embeddings of the Kang dataset. Near and far equidistant cell groups from the activated B cell population were selected following the procedure described in [11]. **(a)** Near equidistant groups in CBMAP (left) and UMAP (right) embeddings. **(b)** Far equidistant groups in CBMAP (left) and UMAP (right) embeddings. Differences in dispersion between near and far groups illustrate how the embedding methods represent relative distance relationships.

A complementary analysis based on the Kolmogorov–Smirnov (K–S) statistic supports the same conclusion (Fig. 7 and Supplementary Fig. S6-S7). This analysis compares the separation between inter-type and intra-type distance distributions. Across datasets, CBMAP generally produced K–S values closer to those observed in the high-dimensional PCA space than the competing embedding methods. By contrast, UMAP, t-SNE, and PaCMAP often increased separation substantially, suggesting that they amplify differences between cell populations beyond what is present in the original data. This is an important point for biological interpretation. Stronger visual separation is not necessarily more faithful. In many cases, it may create the impression of highly discrete populations even when the high-dimensional structure is more gradual or only moderately separated. CBMAP appears more conservative in this regard: it does not maximize visual discreteness but instead preserves separation patterns more consistently with the original data geometry. This behavior is most evident in the Muraro and Utero E8.5 Embryo datasets, where competing methods substantially inflate separation relative to the PCA space. In contrast, in the SMART-seq Mouse dataset, where separation is already high in the reference PCA space, CBMAP produces slightly lower separation values, while the competing methods remain close to or slightly exceed the PCA level.

**Figure 7.**
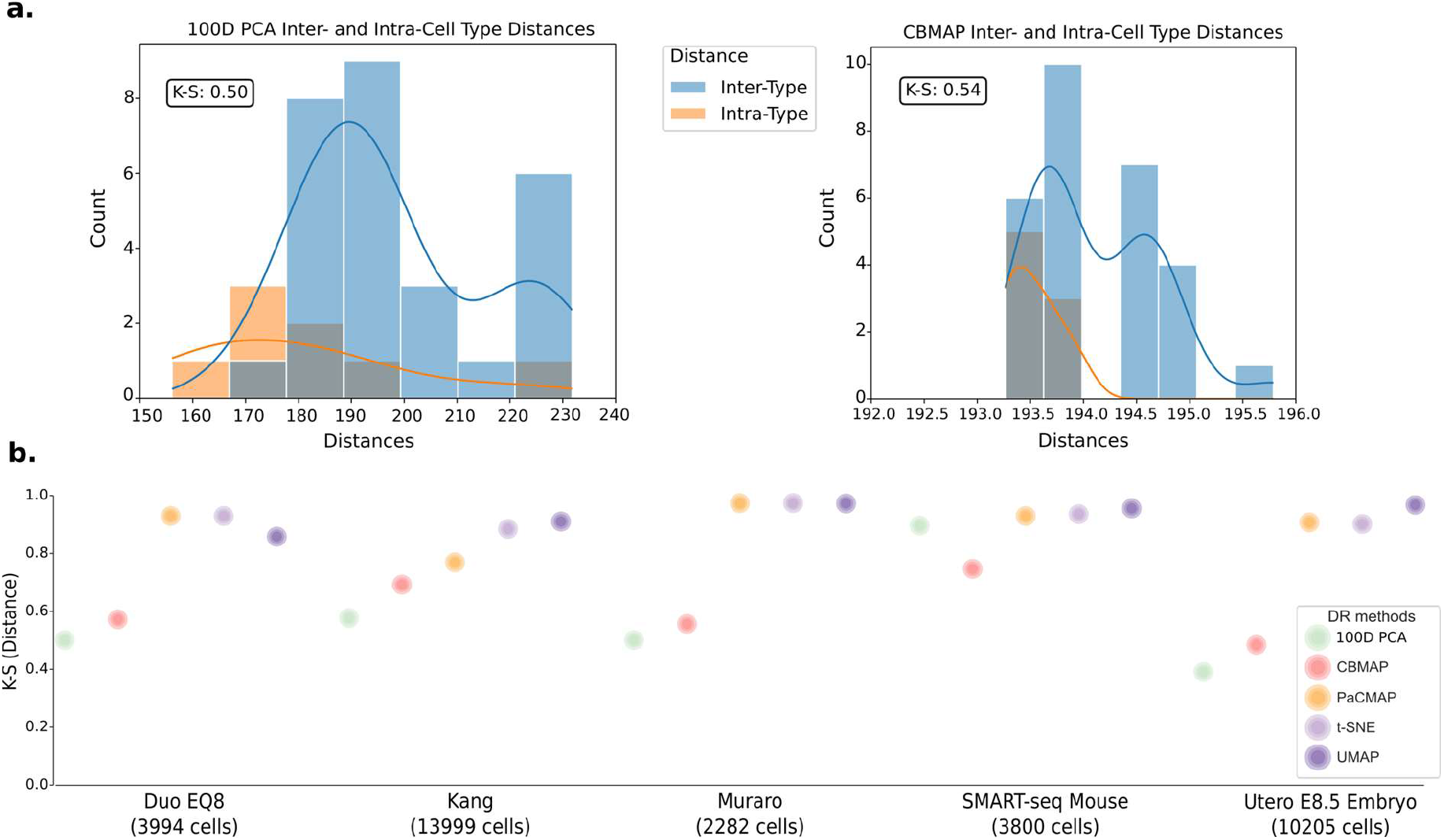
Quantification of cell-type separation using the Kolmogorov–Smirnov (K–S) statistic. **(a)** Distributions of inter-type and intra-type pairwise distances in the high-dimensional PCA space (left) and the corresponding CBMAP embedding (right) for the Duo Eq8 dataset. The K–S statistic summarizes the separation between the two distributions. **(b)** K–S statistics calculated between inter-type and intra-type distance distributions across multiple datasets and dimensionality reduction methods. Higher K–S values indicate greater separation between cell types.

Overall, the results in Figs. 3–7 and Supplementary Figs. S2–S6 indicate that CBMAP provides a more structurally faithful representation of scRNA-seq data. Its main strength is not maximal cluster compactness, but preservation of global organization and relative distance structure, thereby reducing the risk of misleading interpretation caused by visually exaggerated embeddings.

### 3.3. Preservation of developmental trajectories in CBMAP embeddings

To further evaluate the biological interpretability of CBMAP embeddings, we investigated whether the method preserves known developmental trajectories when combined with RNA velocity analysis. For this purpose, CBMAP was evaluated on two well-established trajectory benchmark datasets previously analyzed using the dynamical RNA velocity framework implemented in scVelo: pancreatic endocrinogenesis and dentate gyrus neurogenesis [31–33]. These datasets were selected because their developmental progression and lineage relationships have been extensively validated both computationally and experimentally.

For both benchmarks, we followed the preprocessing, parameter settings, and RNA velocity estimation procedures reported in the reference study [31]. The only modification introduced in our analysis was replacing the original visualization embedding with CBMAP while preserving the same neighborhood graph, velocity vectors, and gene filtering criteria. Clustering was performed using the Secuer algorithm, which showed strong and stable performance in the clustering evaluation stage of this study.

#### Pancreatic endocrinogenesis

The pancreatic endocrinogenesis dataset represents a complex developmental process in which ductal cells give rise to endocrine progenitors that subsequently differentiate into multiple endocrine cell types [31-32]. Previous studies have shown that endocrine progenitors exhibit a characteristic cyclic behavior associated with cell-cycle dynamics before exiting the cycle and committing to specific endocrine lineages.

The CBMAP embedding preserves this biological topology with high clarity (Fig. 8a). The cycling progenitor population forms a coherent loop, and the RNA velocity streamlines trace a smooth transition from this loop toward differentiated endocrine fates. Branching trajectories toward alpha, beta, and other endocrine populations are clearly visible, indicating that CBMAP maintains both cyclic structure and lineage branching in a biologically interpretable manner. Importantly, this is not simply a matter of visual separation; the embedding preserves directional continuity in a way that remains compatible with the inferred developmental flow.

**Figure 8.**
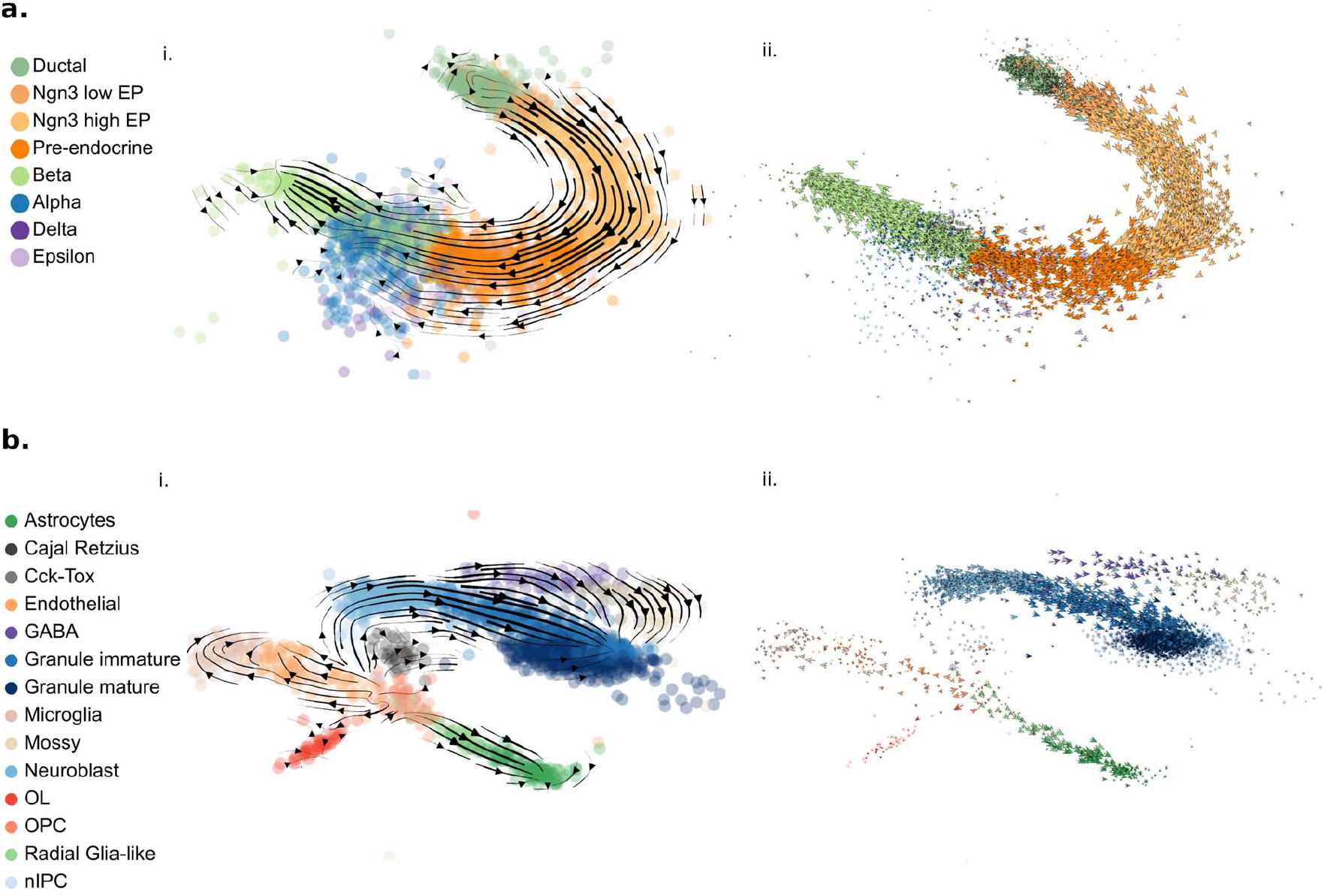
CBMAP visualization of RNA velocity trajectories on developmental benchmark datasets. **(a)** Pancreatic endocrinogenesis. (i) Gene-averaged velocity streamlines (black) projected onto the CBMAP embedding generated using Secuer clustering. (ii) Single-cell velocity vectors (arrows) illustrating local directional transitions from the cycling progenitor loop toward differentiated endocrine states. **(b)** Dentate gyrus neurogenesis. (i) Gene-averaged velocity streamlines (black) projected onto the CBMAP embedding generated using Secuer clustering. (ii) Single-cell velocity vectors highlighting lineage directionality and branching toward terminal cell populations.

#### Dentate gyrus neurogenesis

The dentate gyrus dataset represents a well-characterized neurogenic lineage in the developing brain [31,33]. The main differentiation path proceeds from radial glia to nIPC, neuroblast, granule immature, and granule mature cells, while additional sublineages such as OPC/OL, Cajal–Retzius, and GABAergic populations are also present.

As shown in Fig. 8b, CBMAP preserves this lineage organization with high clarity. The major granule-cell lineage appears as a continuous developmental trajectory aligned with the RNA velocity field, while parallel branches remain distinguishable and biologically interpretable. The single-cell velocity vectors further support this interpretation by showing coherent local directionality along major branches and toward terminal populations. Thus, CBMAP preserves not only large-scale lineage geometry but also local directional consistency relevant to trajectory interpretation.

#### Implications for trajectory-aware visualization

Across both developmental benchmarks, CBMAP maintains the integrity of biologically validated trajectories when combined with RNA velocity information (Figs. 8, S8, and S9). In contrast to UMAP, t-SNE, and PaCMAP, which tend to fragment trajectories into disconnected clusters or distort their global continuity, CBMAP preserves coherent lineage structures and directional flow.

In the pancreatic endocrinogenesis dataset (Fig. S8), CBMAP clearly retains the cyclic progenitor loop and the subsequent branching toward differentiated endocrine states. Competing methods, while producing visually separated clusters, disrupt the continuity of this loop or distort directional transitions, making the trajectory structure less interpretable. Similarly, in the dentate gyrus neurogenesis dataset (Fig. S9), CBMAP preserves the continuous progression of the main lineage while maintaining clear separation of parallel sublineages. In contrast, UMAP and t-SNE tend to fragment the trajectory into isolated components, and PaCMAP partially distorts the global arrangement, limiting the interpretability of lineage relationships.

These observations highlight a key requirement for trajectory-aware visualization: maintaining a balance between global structure and local continuity. CBMAP achieves this balance by preserving lineage-level organization while allowing smooth directional transitions to emerge in the embedding space. Taken together, these results demonstrate that CBMAP is well suited for dynamic trajectory visualization in scRNA-seq data. Its emphasis on global structure preservation enables faithful representation of branching and transitional processes without introducing artificial fragmentation or distortion of biologically meaningful trajectories.

## 4. Conclusion

This study evaluated the applicability of CBMAP as a visualization framework for single-cell RNA sequencing (scRNA-seq) data by systematically examining the impact of different clustering strategies and outlier-aware processing approaches. Multiple clustering algorithms— including k-means, Leiden, HDBSCAN, Secuer, HGC, and FlowSOM—were integrated into the CBMAP pipeline to investigate how clustering behavior affects the structural fidelity of the resulting embeddings. The performance of these configurations was assessed using established global and local structure-preserving metrics and was compared with widely used dimensionality reduction techniques, including UMAP, t-SNE, and PaCMAP.

Our results demonstrate that the clustering stage plays a critical role in the quality of CBMAP visualizations. In particular, outlier-aware clustering strategies substantially improved embedding quality by reducing structural distortions. Across multiple datasets, CBMAP consistently preserved global data organization and inter-population distance relationships more faithfully than the compared methods. Although local neighborhood preservation was generally weaker than in methods explicitly optimized for local structure, the observed distortions were systematic rather than random and remained consistent with the underlying organization of the high-dimensional data.

Importantly, CBMAP embeddings retained biologically meaningful relationships in trajectory-driven datasets. When combined with RNA velocity analysis, CBMAP successfully preserved cyclic progenitor states and branching differentiation trajectories, demonstrating its compatibility with trajectory-aware visualization in single-cell transcriptomic analyses. Beyond its role as a visualization technique, CBMAP also provides a useful diagnostic perspective for clustering evaluation. Because the embedding quality is strongly influenced by clustering behavior, structural distortions observed in CBMAP visualizations can serve as an indirect indicator of clustering performance in scRNA-seq analysis.

Future work should focus on improving the CBMAP membership formulation to better accommodate the sparse and heterogeneous characteristics of single-cell transcriptomic data. Such developments may further enhance local structure preservation while maintaining the strong global structure fidelity observed in this study.

## Data and code availability

The code and datasets supporting the findings of this study are publicly available in the GitHub repository: https://github.com/doganlab/cbmap-scRNAseq

The repository includes all scripts required to reproduce the analyses, generate the results, and perform the benchmarking experiments presented in this work. Further information is available from the corresponding author upon reasonable request.

## CRediT authorship contribution statement

**Meysa Alchaar:** Software, Formal analysis, Visualization, Writing – review & editing. **Berat Doğan:** Conceptualization, Methodology, Writing - original draft.

## Notes

### Competing Interest Statement

The authors have declared no competing interest.

