## Supplementary Figures for "Clustering Strategies Improve Structure-Preserving Visualization of Single-Cell RNA-seq Data with CBMAP"

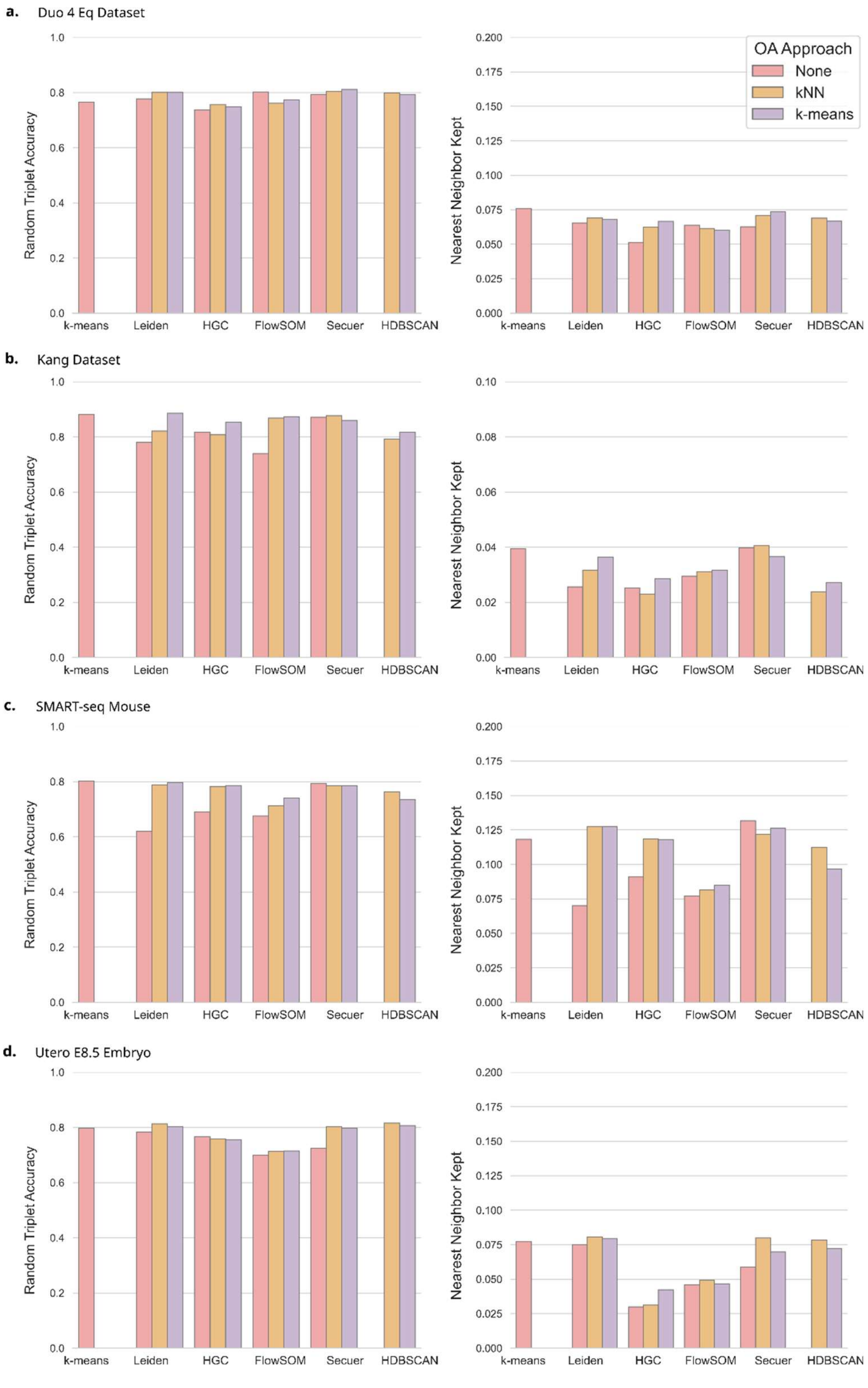


**Figure S1**. CBMAP embeddings obtained using different clustering approaches across multiple scRNA-seq datasets. Bar plots represent the average evaluation metrics across five runs for each clustering algorithm. Default parameters were used unless otherwise specified. Three outlier-handling strategies were considered: no outlier handling, sigma-rule–based outlier detection followed by assignment to the nearest inlier cluster using k-nearest neighbors (kNN), and clustering of detected outliers using k-means. **(a)** Duo4Eq dataset: k-means, HGC, and FlowSOM were run with 24 clusters (three times the true number of cell types); Leiden resolution was set to 0.5; HGC used 20 nearest neighbors. **(b)** Kang dataset: k-means used 39 clusters; HGC and FlowSOM used 13 clusters; Leiden resolution was set to 2.5; HGC used 15 nearest neighbors. **(c)** SMART-seq Mouse dataset: k-means used 40 clusters; HGC and FlowSOM used 28 clusters (equal to the number of annotated cell types); Leiden resolution was set to 0.5; HGC used 30 nearest neighbors. **(d)** Utero E8.5 Embryo dataset: k-means used 56 clusters; HGC and FlowSOM used 14 clusters (true number of cell types: 19); Leiden resolution was set to 0.7; HGC used 20 nearest neighbors. For all panels, global structure preservation (right) is evaluated using the Random Triplet Accuracy (RTA) metric, and local structure preservation (left) is evaluated using the Nearest Neighbor Kept (NNKept) metric.

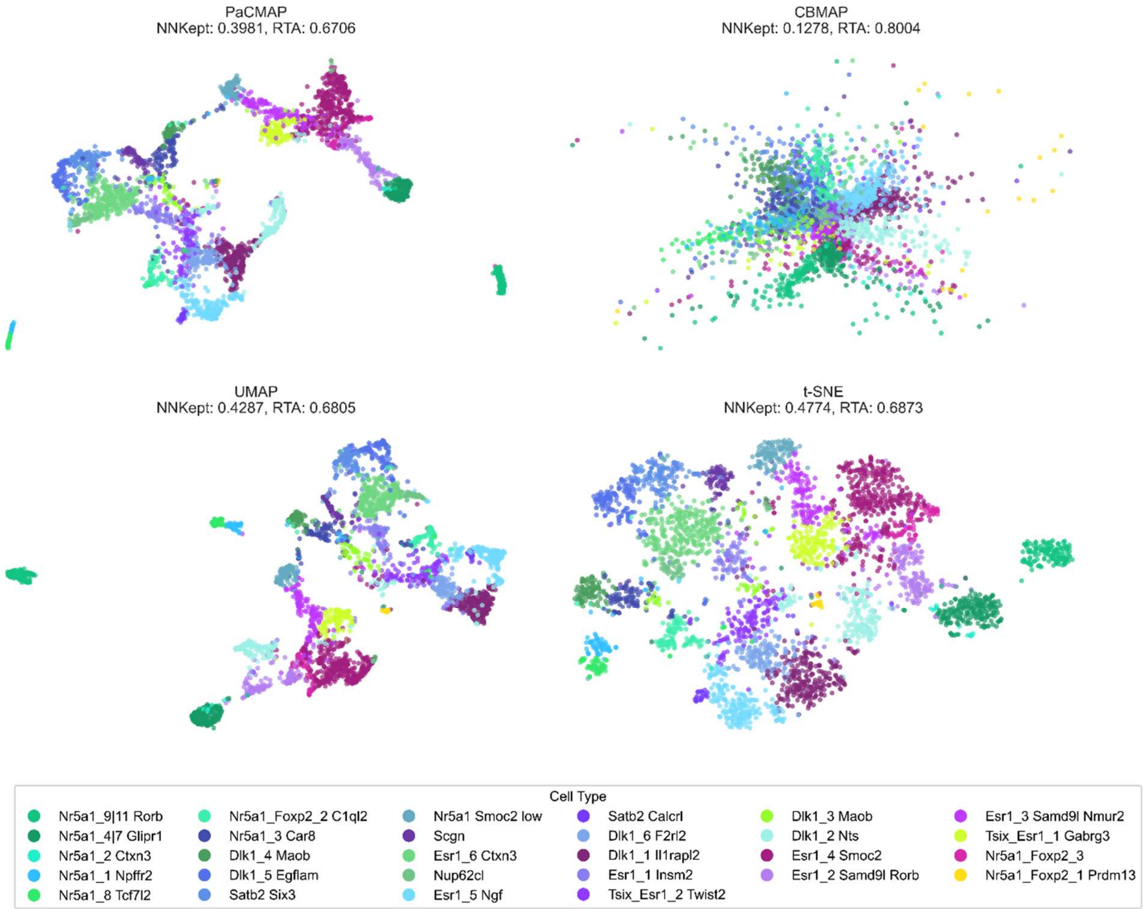


**Figure S2.** Comparison of different embedding techniques on the SMART-seq Mouse dataset. All embedding methods were implemented using default parameter settings. For CBMAP, the Leiden clustering algorithm (resolution: 0.5, OA-approach: kmeans) was used to generate the embeddings.

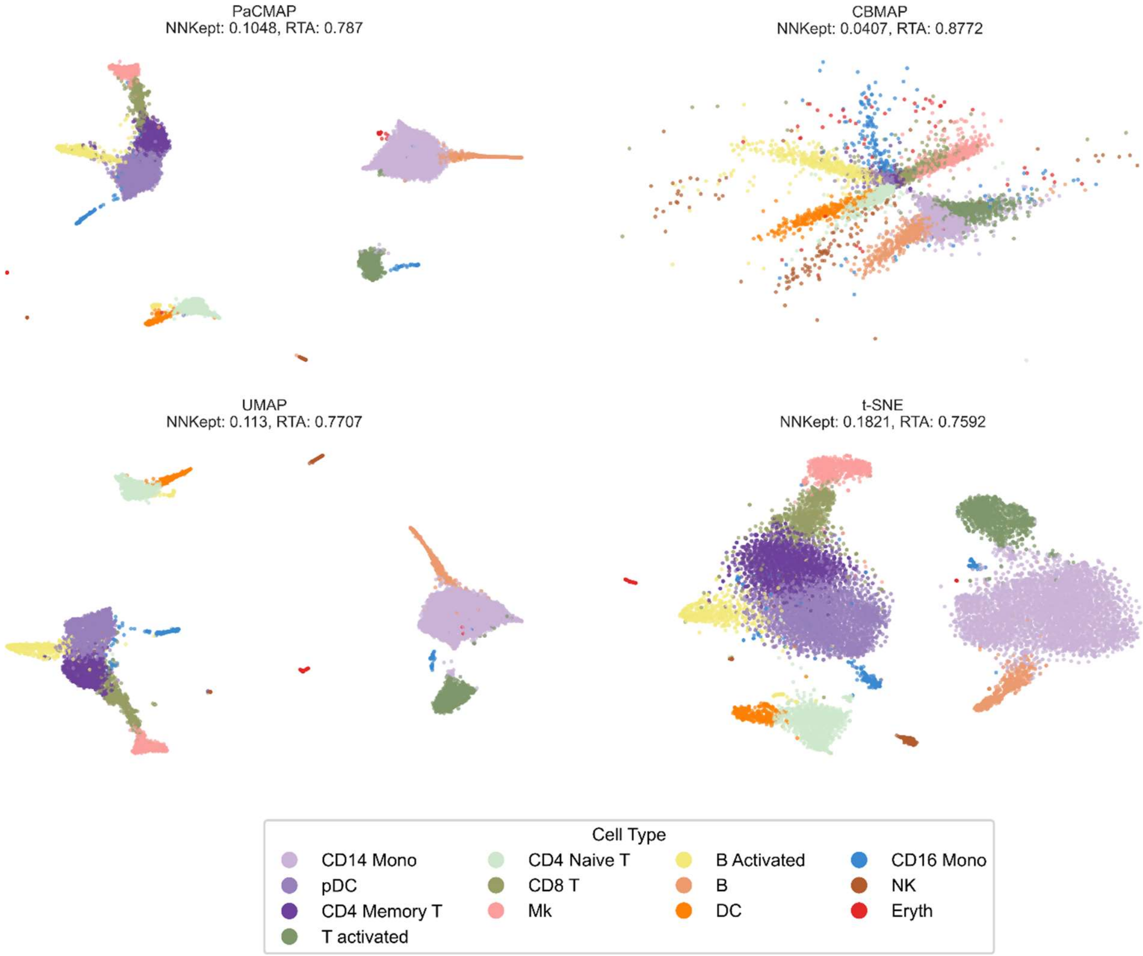


**Figure S3.** Comparison of different embedding techniques on the Kang dataset. All embedding methods were implemented using default parameter settings. For CBMAP, the Secuer clustering algorithm (OA-approach: kNN) was used to generate the embeddings.

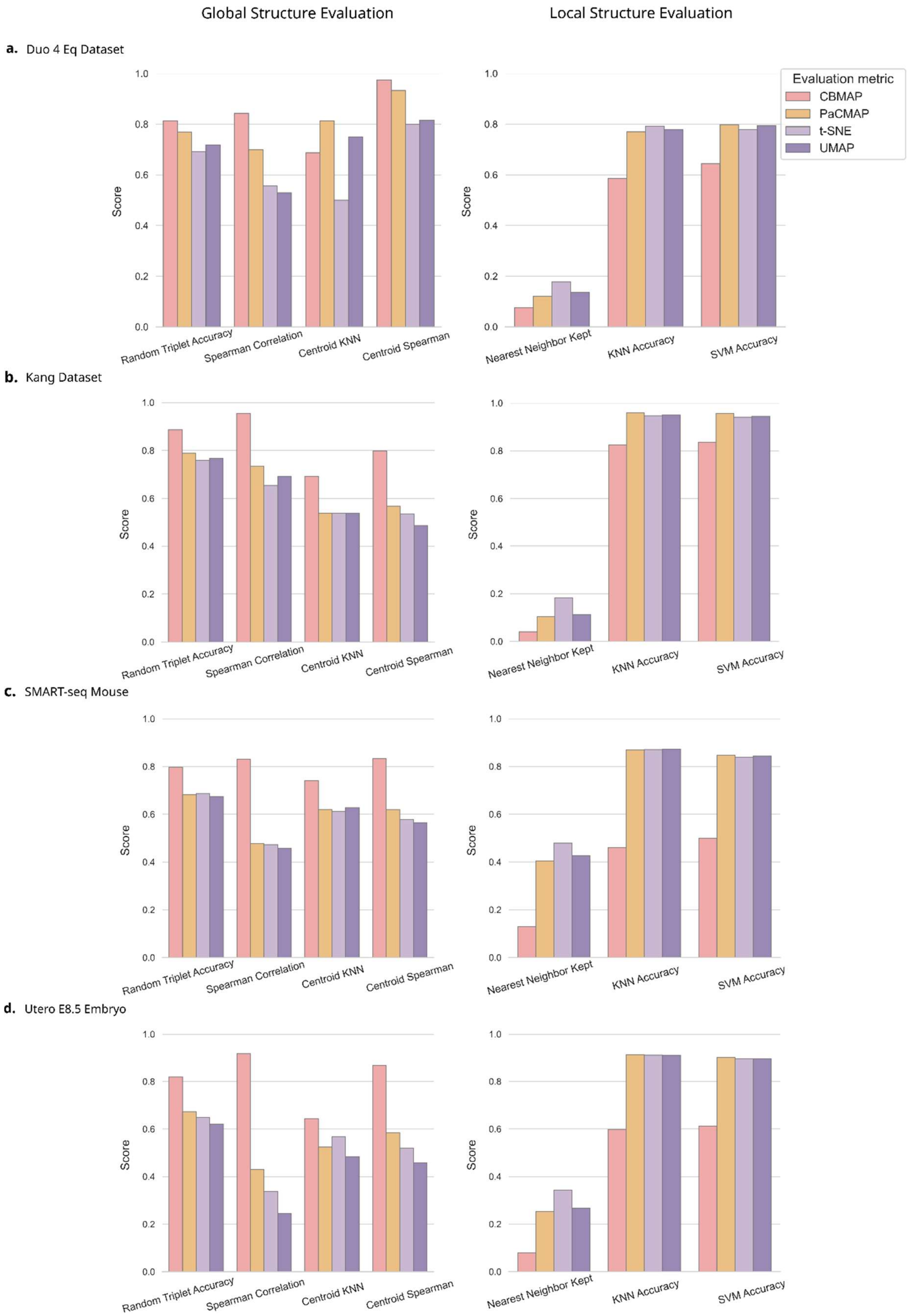


**Figure S4.** Evaluation of structure preservation for different embedding techniques. Bar plots represents the average evaluation metrics across five runs for each dimensionality reduction method. (Left) Global structure preservation metrics (Right) Local structure preservation metrics **(a)** Evaluation of structure preservation for different embedding techniques on the Duo 4 Eq dataset. **(b)** Evaluation of structure preservation for different embedding techniques on the Kang dataset. **(c)** Evaluation of structure preservation for different embedding techniques on the SMART-seq Mouse. **(d)** Evaluation of structure preservation for different embedding techniques on the Utero E8.5 Embryo dataset.

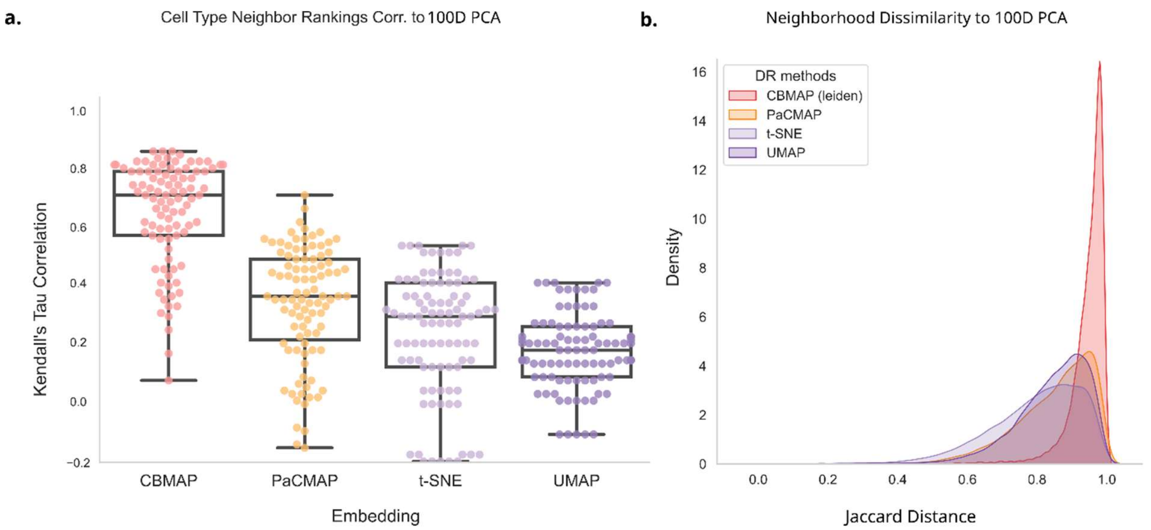


**Figure S5.** Evaluation of structure preservation for different embedding techniques on the Utero E8.5 Embryo dataset. **(a)** Boxplots showing the correlations between cell-type neighbor rankings in the 2D embeddings and the corresponding higher-dimensional PCA space. Each embedding method was generated five times; higher correlations indicate better structure preservation. **(b)** Jaccard distance distribution of cell neighbors in 2D embedding spaces relative to the higher-dimensional PCA space; lower Jaccard distances indicate better agreement.

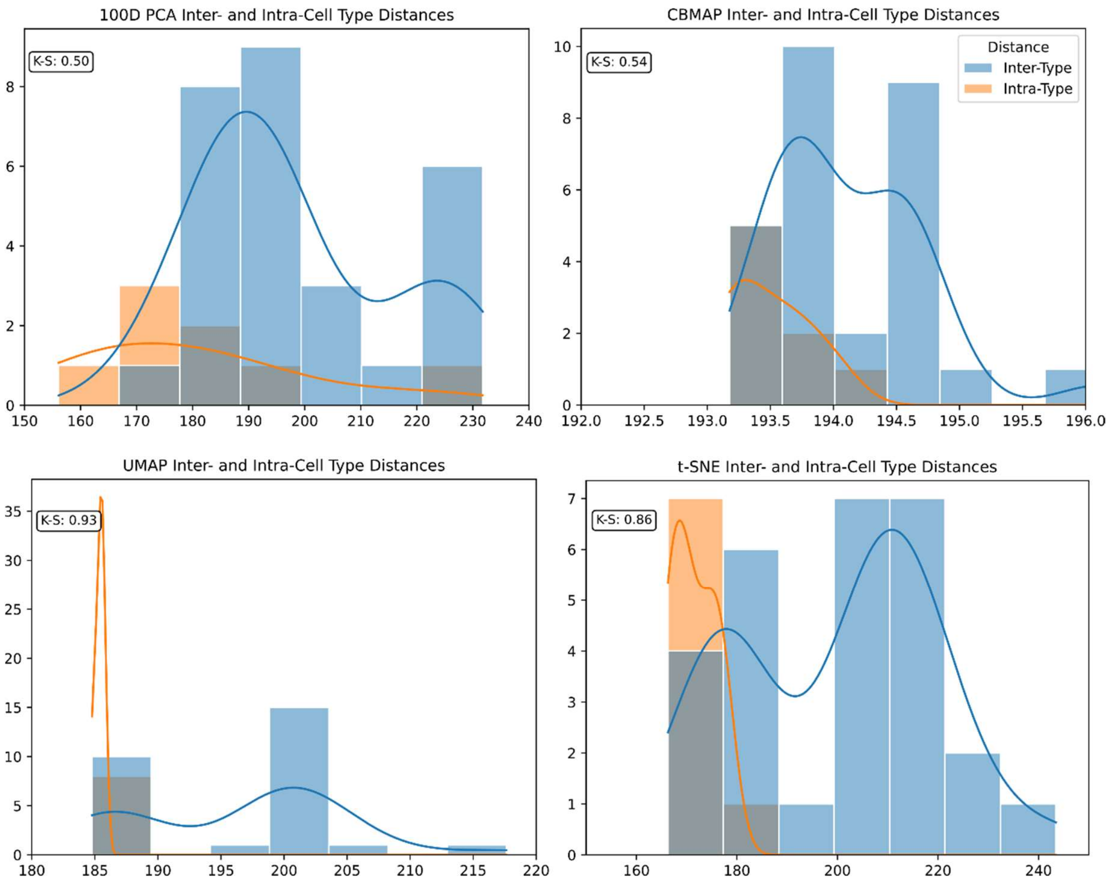


**Figure S6.** Quantifying cell-type separation using the Kolmogorov–Smirnov (K–S) statistic for the Duo Eq8 dataset. Distributions of inter- and intra-type pairwise distances in the high-dimensional PCA space and in the corresponding embedding methods of the Duo Eq8 dataset. The K–S` statistic summarizes the degree of separation between the two distributions.

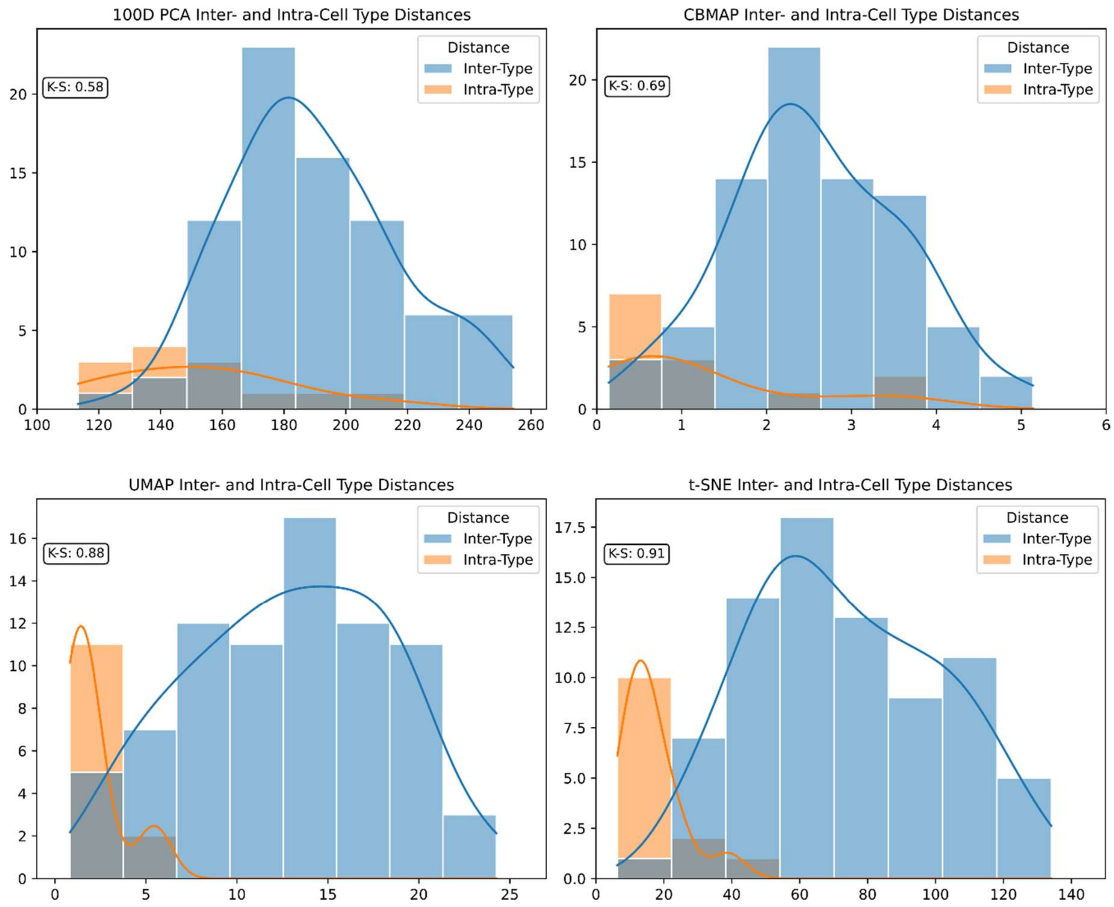


**Figure S7.** Quantifying cell-type separation using the Kolmogorov–Smirnov (K–S) statistic for Kang dataset. Distributions of inter- and intra-type pairwise distances in the high-dimensional PCA space and in the corresponding embedding methods of the Kang dataset. The K–S` statistic summarizes the degree of separation between the two distributions.

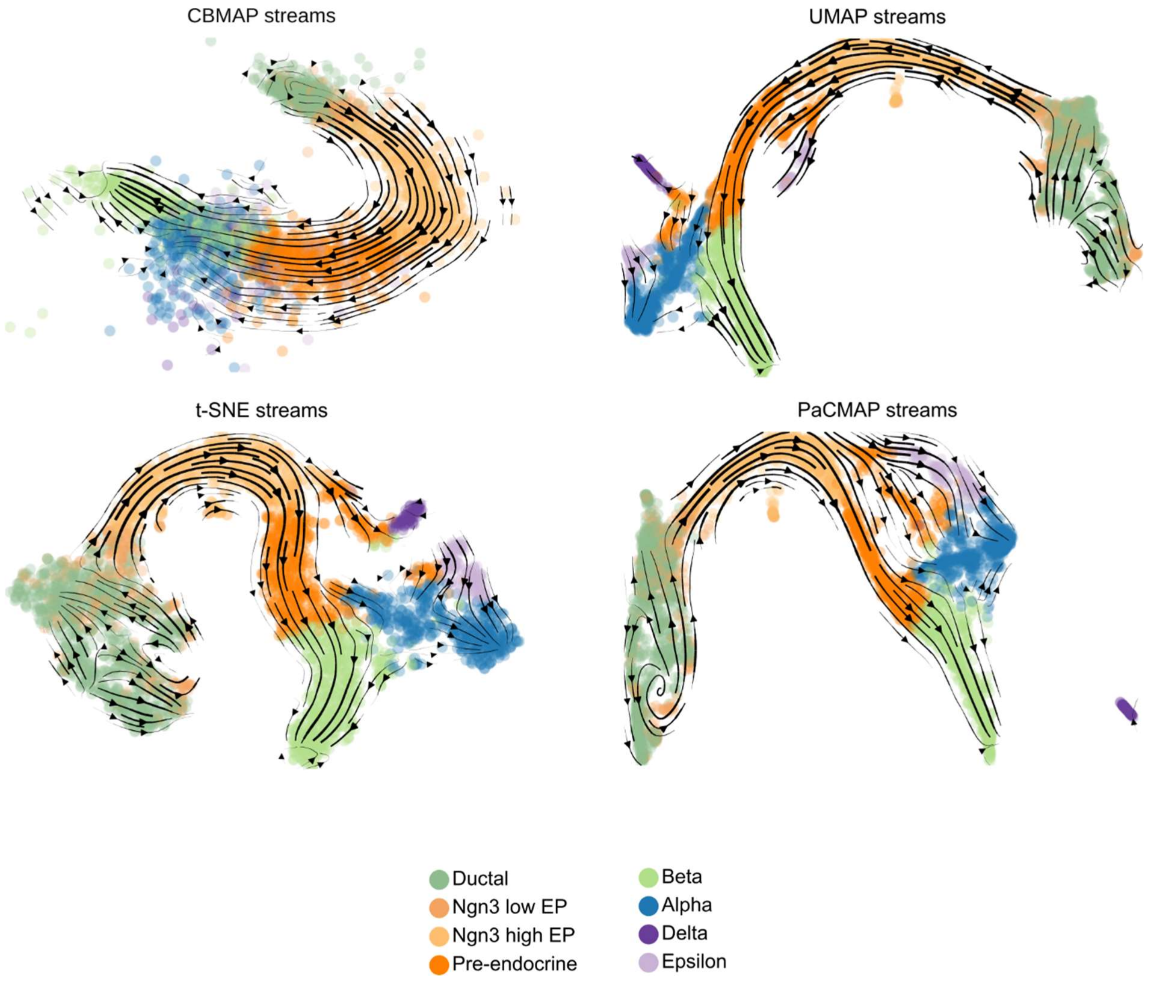


**Figure S8.** Comparison of RNA velocity trajectory projections across 2D embedding methods in pancreatic endocrinogenesis. Gene-averaged RNA velocity streamlines (black) are overlaid on embeddings generated using CBMAP, UMAP, t-SNE, and PaCMAP. All embeddings are computed from the same top 30 principal components (PCA space) using default parameter settings. The CBMAP embedding is generated using the Secuer clustering algorithm.

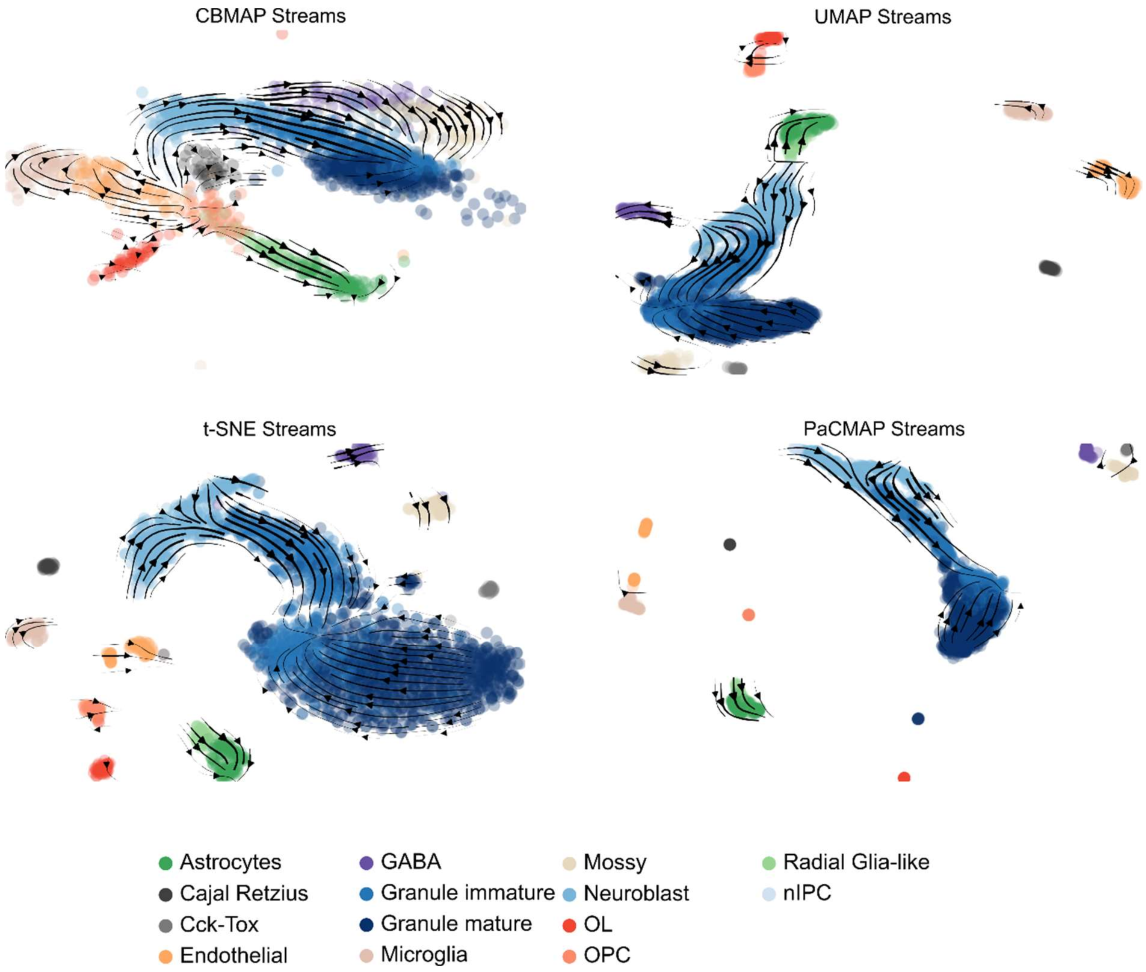


**Figure S9.** Comparison of RNA velocity trajectory projections across 2D embedding methods in dentate gyrus neurogenesis. Gene-averaged RNA velocity streamlines (black) are overlaid on embeddings generated using CBMAP, UMAP, t-SNE, and PaCMAP. All embeddings are computed from the same top 30 principal components (PCA space) under default parameter settings. For UMAP, the number of neighbors is set to 30, consistent with the reference study. The CBMAP embedding is constructed using the Secuer clustering algorithm.
